# The Application of 3D Cryo-Volume Electron Microscopy to Visualize and Quantify the Photosynthetic Filamentous Cyanobacterium *Anabaena*

**DOI:** 10.64898/2026.11.02.763640

**Authors:** Allyson Angermeier, Lolita G. Rotkina, Clara Lebow, Wah Chiu, Tessa M. Burch-Smith, Kirk J. Czymmek

**Affiliations:** Donald Danforth Plant Science Center, Saint Louis, MO, United States; Donald Danforth Plant Science Center, Advanced Bioimaging Laboratory, Saint Louis, MO, United States; Stanford University–Stanford CA, United States

**Author notes:** Corresponding author: Kirk J. Czymmek, Advanced Bioimaging Laboratory, Donald Danforth Plant Science Center, 975 North Warson Road, Saint Louis, MO 63132, United States.

**Keywords:** cryogenic volume electron microscopy, plasma focused ion beam, *Anabaena*, carboxysome, cyanophycin granules, phycobiliomes, thylakoid membranes, correlative cryo-fluorescence

## Abstract

Cryogenic volume electron microscopy (cryo-vEM) is a rapidly developing nanoscale imaging approach that enables three-dimensional datasets of vitrified cells and tissues at intermediate resolutions without treatment with chemical fixatives or metal contrasting agents. Our goal was to evaluate the suitability of cryo-vEM for whole cells of photosynthetic organisms such as cyanobacteria. Two species of *Anabaena*, a filamentous cyanobacterium, were captured on poly-L-lysine-coated grids, vitrified by plunge-freezing into liquid ethane, and mapped by secondary electron and integrated fluorescence light microscopy to identify target cells. Cells of interest were coated with platinum, trenched with xenon, and 3D volumes collected by alternating argon milling and through-lens detector (TLD) imaging on a Thermo Scientific Hydra plasma focused ion beam scanning electron microscope. Volumes were segmented via 2.5D U-Net models, then quantified in Dragonfly 3D World. Block-face images revealed the cell wall, thylakoids, phycobilisomes, carboxysomes, cyanophycin and polyphosphate granules. Whole-cell 3D rendering allowed quantification of carboxysomes, cyanophycin granules, and measurement of thylakoid thickness and spacing in *A. inaequalis* and *Anabaena* sp. UTEX 2576. Thus, cryo-vEM provided interpretable whole-cell ultrastructure of cyanobacteria without chemical pre-treatment, and its compatibility with cryo-fluorescence and *in situ* cryo-ET enables contextual multi-scale perspectives from whole cells to molecular structures.

## 1 Introduction

Since its inception, electron microscopy (EM) has enabled the study and interpretation of cellular structures, organelles, and proteins at the nanoscale. Sample preparation for EM via conventional chemical fixation typically includes aldehydes, osmium tetroxide, heavy metal staining, and embedding samples in resin. Classical EM processing methods are time consuming, often taking 2 weeks, are sample dependent, and come with fixation artifacts depending on the biological question and approach. Additionally, while conventional fixation and staining with heavy metals certainly increases contrast within the sample, in some instances it can significantly alter or even destroy certain cellular structures. Furthermore, aldehyde cross-linking is slow, so structures can move, aggregate, or redistribute before they are immobilized [1], and solvent dehydration and resin infiltration can induce non-linear shrinkage and extract cellular membranes, lipids and proteins [2]. To overcome many of the limitations of conventional chemical fixation and view cellular structures of whole single cells in their native hydrated state, cryogenic EM (cryo-EM) was applied.

Cryo-EM techniques depend on uniform vitrification of a biological specimen [3]. Vitrification is a relatively quick process, immobilizing the sample in milliseconds as opposed to minutes or longer using conventional fixation [4]. Since successful vitrification can more rapidly preserve cellular structures in their native states without requiring exogenous chemicals, the integrity of some biological phenomena (e.g., labile or dynamic structures, spatial relationships) are better preserved. While the cooling rate decreases rapidly with thickness [5], plunge-freezing on EM grids is limited to aqueous layers of a few micrometers [6], suitable for many cyanobacteria. High-pressure freezing extends vitrification to specimens up to approximately 200 µm (sample dependent), which is more appropriate for bulk samples and multicellular tissues [7] or concentrated single cells [8]. The Marko lab conducted the first demonstrated application of focused ion beam (FIB) for frozen hydrated specimens [9], and subsequently the Villa lab achieved lamellae milled from intact vitrified cells [10], enabling intact cellular cryo-ET and ultimately to molecular resolution imaging *in situ* [11]. While *in situ* cryo-ET of lamellae has provided remarkable perspectives of macromolecular detail in cellular context, lamellae are restricted to 100–300 nm thick and only a small fraction of any cell can be captured.

Building upon visualizing the native structures of biological samples, cryogenic volume electron microscopy (cryo-vEM) has more recently gained attention. The cryo-vEM method utilizes the focused ion beam on an SEM to mill a thin slice of the specimen, then alternates to the electron beam to acquire an image of the freshly exposed block face. This process can be repeated until a 3D volume of the sample is acquired. The approach was first demonstrated by Schertel et al. on high-pressure frozen, unstained mouse optic nerve and *Bacillus subtilis* spores [12], then extended to large volumes of frozen hydrated tissue [13], to whole vitrified cells within a correlative multi-scale pipeline [14], and beyond. Without the introduction of any heavy metals, contrast is not fully understood, but arises from the specimen itself [12, 15]. The approach is compatible with gallium ion or an inductively coupled plasma source and cryogenic plasma FIB-SEM (cryo-pFIB-SEM) which allows serial block-face imaging of larger volumes [16]. Nevertheless, cryo-vEM is not without its challenges, including local charging, (typically requiring sample and system dependent optimization) [17], beam damage, and some cell features may not be well contrasted.

Since cryo-vEM volume acquires small regions relative to the grid or bulk samples, for efficiency, knowing where to mill is important. For this, correlative cryo-fluorescence has become a reliable solution and with developing three-dimensional (3D) correlation approaches it is possible to establish a position and even depth of a target within the cell in advance of milling [18] and an integrated fluorescence microscope inside the FIB-SEM chamber is convenient for this task [19, 20]. Fluorescence has also been used to guide a volume acquisition directly rather than simply to position a lamella [14, 21]. After acquisition, cryo-vEM data can be imported into analysis software such as DeepMIB [22, 23], Napari [24], Empanada [25] or commercial solutions such as Dragonfly 3D, ARIVIS or Amira, for visualization, qualitative and quantitative analyses. Within these tools, features of interest can be segmented manually or with the assistance of deep learning AI models [26], which becomes increasingly important with while cells and 100s or 1000s of slices [27].

While development of these cryogenic techniques thus far has mostly been applied to unicellular organisms or mammalian cells and tissues, photosynthetic organisms have been limited. However, cryo-FIB-SEM milling combined with cryo-ET was applied to determine architecture of the *Chlamydomonas* chloroplast and pyrenoid by Engel et al. [28], thylakoid organization and plasma membrane contact sites in *Synechocystis* by Rast et al. [29], and Weiss et al. characterized the septal junctions between cells within *Anabaena* filaments [30]. Volume EM of plant cells in general has relied on chemical fixation, or on high-pressure freezing followed by freeze-substitution and heavy-metal staining [31, 32, 33], so that the specimen is chemically modified before imaging. True volume cryo-vEM of photosynthetic cells, by contrast, is limited to a handful of reports, including a coccolithophorid alga [34], whole-cell *Chlamydomonas* [35], guard cells of *Vicia faba* [36], and sorghum protoplasts [37]. Cryogenic approaches are only beginning to reach plant material, so for a good starting point for multicellular tissue combined with FIB-SEM for *in situ* cryo-ET workflows, see Poge et al. [38]. To our knowledge, cryo-vEM has not previously been applied to a cyanobacterium.

Our long-term goal is to extend the benefits of cryo-vEM for 3D imaging of whole cells of a variety of photosynthetic organisms, including multicellular plants and tissues, and to build upon the work of others. Here we focus on *Anabaena*, a filamentous cyanobacterium whose biology, heterocyst differentiation, intercellular transport through septal junctions, and carbon fixation within carboxysomes [39, 40], is organized at the mesoscale that falls between the geometry and resolution of cryo-ET lamella and light microscopy. We describe the complete workflow from vitrification through cryo-pFIB-SEM acquisition, segmentation and volumetric quantification, and consider what native-state imaging adds for structures such as carboxysomes, thylakoid membranes, polyphosphate granules, and cyanophycin granules that are prone to extraction, low contrast or alteration during conventional preparation [41, 42]. In general, multicellular plants can be more challenging for cryogenic imaging due to their large central vacuoles, cell walls, waxy cuticles, and air spaces that impede space fillers and vitrification, especially in multi-cellular leaf tissues. Nevertheless, as cryo-vEM becomes more accessible, there is a very strong need to develop robust routines in photosynthetic organisms that leverage the benefits of imaging vitrified frozen hydrated specimens without chemical pretreatment.

## 2 Materials and methods

### 2.1 Filamentous cyanobacteria culture

*Anabaena* sp. UTEX 2576 (Culture Collection of Algae at the University of Texas at Austin (UTEX)) and *Anabaena inaequalis* (Carolina Biological Sciences Item #151710) cultures were grown in BG-11 medium (UTEX, https://utex.org/products/bg-11-medium?variant=30991786868826) at 27°C on a 12:12 hour light/dark cycle. Cultures were grown in a photobioreactor (UTEX Photobioreactors; Plus Package with RGB-LED Lighting Platform) and supplied with atmospheric CO_2_ via an air pump. Immediately prior to freezing, culture density was measured using OD_750_ in a VWR® UV-1600PC spectrophotometer to ensure they were in log phase growth.

### 2.2 Grid preparation

UltrAuFoil® 300 mesh Au grids (Cat.# 688-300-AU-50, Tedd Pella, California, US) were glow discharged in a Leica EM ACE600 sputter coater (Leica Microsystems, Vienna, Austria) for 2 minutes. In order to improve consistency in filament attachment, the grids were coated with 0.1% Poly-L-Lysine (Cat.# P4707, Millipore Sigma, Massachusetts, US) for 1 minute, blotted at the edge with filter paper to remove excess liquid, then allowed to dry on a grid prep holder (Cat.# 71175-00, Electron Microscopy Sciences, Pennsylvania, US) inside a covered petri dish overnight at room temperature and summarized in **Figure 1** (Grid Preparation).

**Figure 1.**
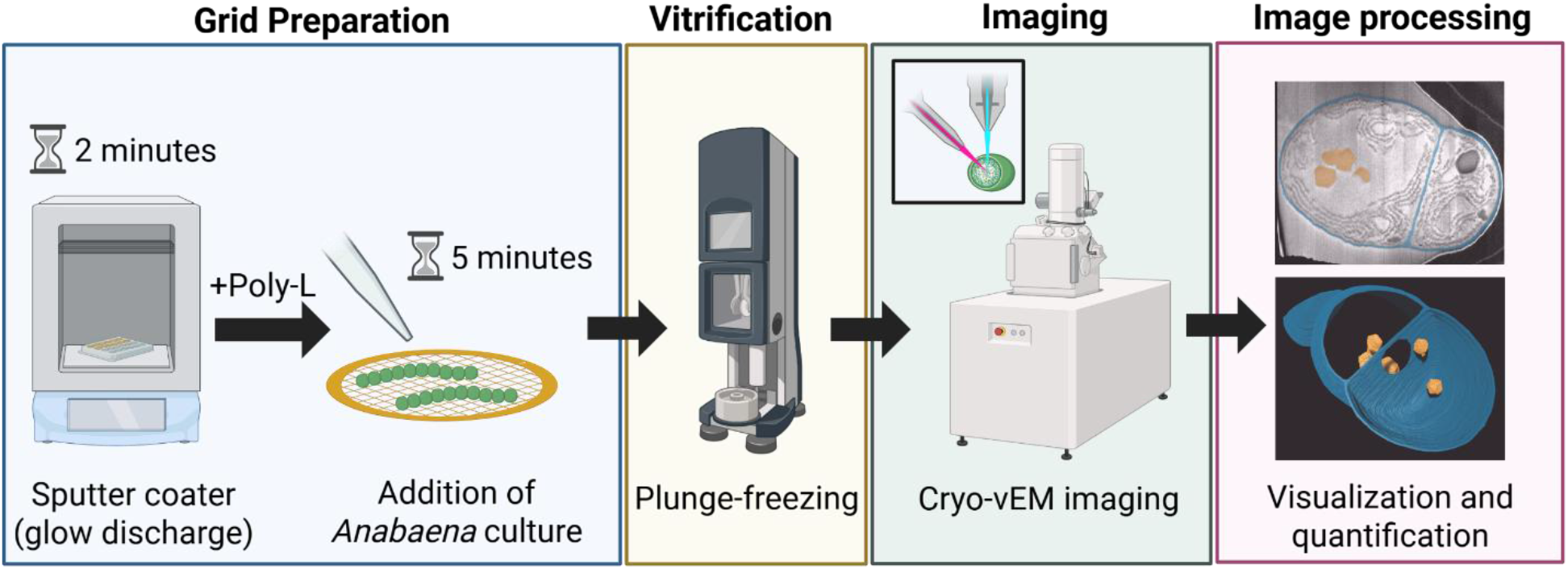
Cryo-vEM workflow. *Grid Preparation* - UltrAuFoil 300 mesh grids were initially prepared by using an argon beam in a Leica EM ACE600 to displace hydrophobic ions on the grid to make them more hydrophilic (glow discharging), then treated with aqueous poly-L-lysine (poly-L). After drying the poly-L-coated grids overnight, *Anabaena* culture was added to the grids for 5 minutes. *Vitrification* - Grids were transferred to a Leica EM GP2 and plunge-frozen. I*maging* - After vitrification, the grids are placed into the Helios Hydra cryo-pFIB-SEM equipped with iFLM for targeting and 3D cryo-vEM imaging. *Image processing* -The resulting image stacks were annotated, segmented, quantified and visualized in the Dragonfly 3D World.

### 2.3 Plunge-freezing

The cryo-chamber of the Leica EM GP2 plunge freezer (Leica Microsystems, Vienna, Austria) was pre-cooled using liquid nitrogen at least 15 minutes before use. The cryo-chamber remains chilled by vapor phase liquid nitrogen. Ethane gas was added to a small cup placed beneath the plunger within the cryo-chamber, where it condensed into liquid ethane upon cooling by liquid nitrogen. A second small insert was filled with liquid nitrogen where a hybrid grid holder (Cat.# 14021008, Nanosoft, Rhode Island, US) is placed for storage and transfer of vitrified grids. A piece of blotting paper supplied by Leica was placed into the blotting chamber. Grids were held using Leica EM GP2 forceps and placed horizontally on the bench, poly-L-lysine coated side up. Then, an 8 µL droplet of *Anabaena* culture was placed onto each grid for 5 minutes at room temperature to allow cell capture onto the poly-L-lysine coated surface before vitrification. After 5 minutes, the forceps and grid were loaded into the pre-cooled plunge freezer (**Figure 2E–G**). Samples were then automatically loaded into the Leica EM GP2 plunge freezer chamber at 41% humidity (ambient) and single blotted using the blot sensor feature. The blotting time was 0.5 seconds with an additional move of 2.1 mm before being plunged into liquid ethane for vitrification (blotting conditions need to be empirically determined for your sample and system).

**Figure 2.**
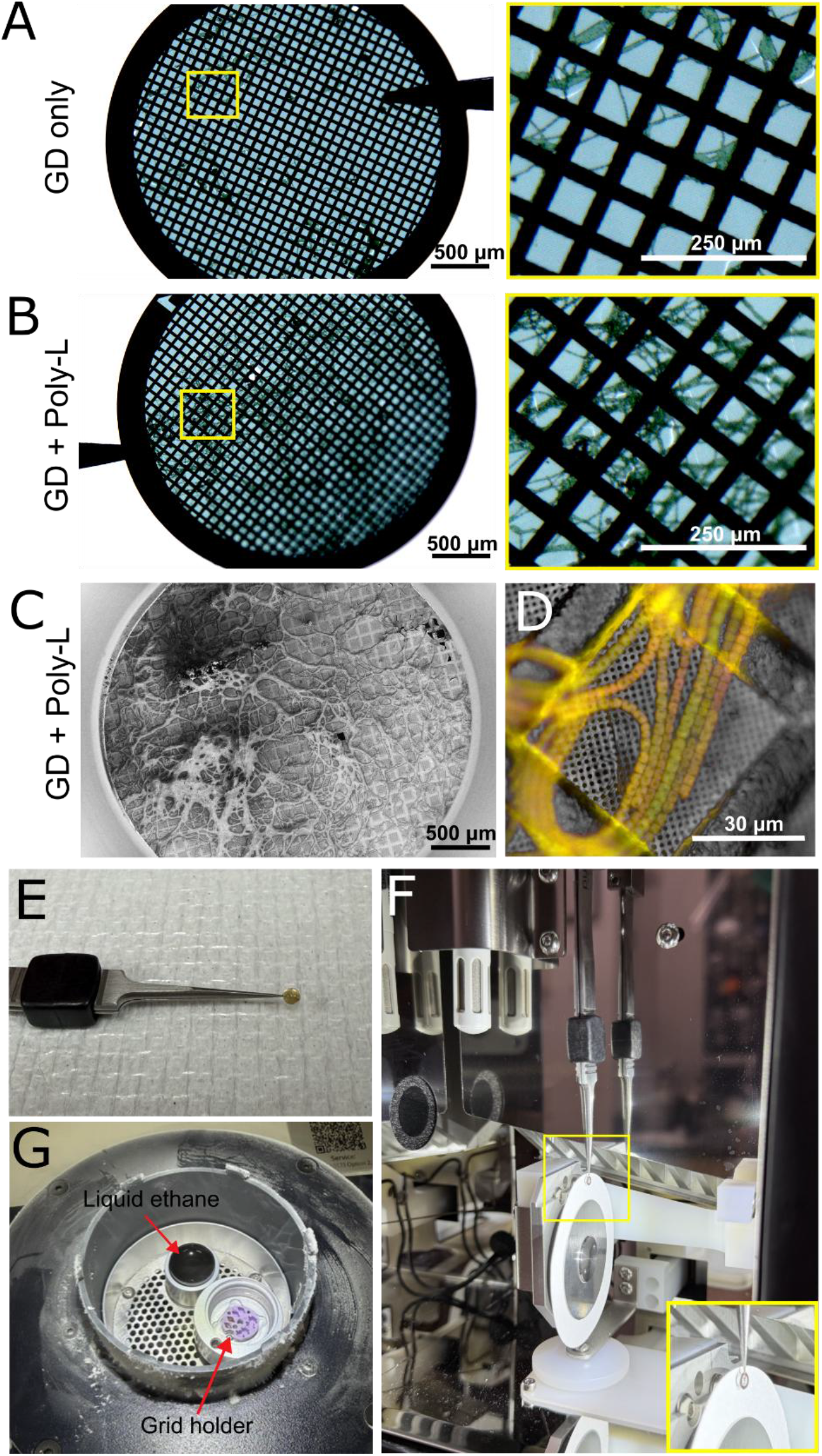
Poly-L-lysine treatment and vitrification via plunge-freezing. (A) Brightfield image of *Anabaena* on a grid after glow discharge and blotting in the plunge freezer at room temperature (left) and zoomed (right). (B) Brightfield image of *Anabaena* on a grid after glow discharge, poly-L-coating, and blotting in the plunge freezer at room temperature (left) and zoomed (right). (C) FIB-SEM map of *Anabaena* on a grid that was glow discharged, poly-L-coated, and vitrified in the plunge freezer. (D) A zoomed-in image of *Anabaena* on a grid detected using iFLM. (E) An image of the *Anabaena* incubation for 5 minutes. The grid is held in forceps and 8 µL of *Anabaena* culture in liquid media is added to the grid and allowed to adhere. (F) After the *Anabaena* incubation, the forceps with grid are introduced into the plunge-freezer chamber to be blotted with the *Anabaena* culture in the direction of the blot paper. (G) The cryo-chamber of the plunge freezer showing the ethane liquid as well as the grid holder with two vitrified grids stored inside.

After vitrification, the forcep holding the grid was released from the plunge-freezer and quickly transferred within the plunge freezer cryo-chamber to a hybrid grid box (Cat.# 14021008, Nanosoft, Rhode Island, US). A pin lid was attached to the grid box to contain the grids using a pin lid manipulator tool (Cat.# 27011001, Nanosoft, Rhode Island, US). After the grids were secured in the grid box, they were carefully transferred to a MiTeGen autogrid storage puck (Cat.# 28021004, Nanosoft, Rhode Island, US) and stored in a liquid nitrogen dewar until clipping and subsequent imaging. All steps were performed to minimize exposure to ambient air and ice contamination.

### 2.4 pFIB-SEM imaging for cryo-vEM

Prior to imaging, the vitrified grids were clipped in a grid clipping station (Cat.# 71167-90, Electron Microscopy Sciences, Pennsylvania, US) filled with liquid nitrogen. The vitrified grids were removed from storage and placed into the clipping container. A C-clip (Cat.# 71167-97, Electron Microscopy Sciences, Pennsylvania, US) was placed into clipping tool (Cat.# 71167-94, Electron Microscopy Sciences) and around the grid to create an AutoGrid assembly (Cat.# 71167-97, Electron Microscopy Sciences). AutoGrids were mounted on an Aquilos 35° pre-tilted cryo-shuttle holder and transferred to the cryo-stage cooled to -170°C. A first conductive platinum layer was applied by micro-sputter coating (xenon, 12 kV, 0.12 µA, micro-sputterer pattern, 120 s). Serial FIB-SEM volume imaging was performed on a Thermo Fisher Scientific Helios 5 Hydra CX plasma FIB-SEM (Thermo Fisher Scientific, Massachusetts, US) equipped with four plasma sources (xenon, argon, oxygen and nitrogen), an integrated fluorescence light microscope (iFLM) and a cryo-stage, using the Auto Slice & View (ASV, Version 5.10) automated serial mill-and-image workflow for cryogenic specimens.

Grids were then mapped in MAPS software (Version 3.24 1.33, Thermo Fisher Scientific, Massachusetts, US) and candidate *Anabaena* filaments were located by correlative fluorescence imaging prior to milling. Because *Anabaena* grows as long chains of cells and its chlorophyll and other cyanobacteria pigments have strong, broad spectrum autofluorescence, no exogenous labelling was required: fluorescence map z-stacks were acquired for candidate *Anabaena* filaments using green (470nm) and red (565nm) excitation and accordant emission channels appropriate to the intrinsic autofluorescence of the specimen. The MAPS and iFLM stacks were aligned to one another for target selection as needed.

The protective multilayer coating was then completed with four further deposition steps: platinum deposition from the gas injection system (GIS) at 27 °C for 30–45 s (layer 2); a second xenon micro-sputter coating (12 kV, 0.12 µA, 120 s; layer 3); a third GIS platinum deposition for 30–45 s (layer 4); and a final xenon micro-sputter coating (12 kV, 0.12 µA, 360 s; layer 5).

For the data presented herein, two whole cell volumes were acquired (N = 1 complete cell for each species, partial cells not included herein) - one approaching the filament from the side (longitudinal axis) and one as an oblique cross-section to the filament; both were milled and imaged at the same stage position, with the specimen surface perpendicular to the electron beam. Targeted sites were trenched and fiducials milled using the xenon plasma source at 30 kV, with beam current selected according to the precision required at each step. The plasma source was then switched to argon for final site preparation and for automated serial milling. For acquisition, the stage was tilted to 35° at a working distance of 4.0 mm so that the grid surface lay perpendicular to the electron beam, giving a FIB milling angle of 52° relative to the specimen surface. A fiducial pattern was milled above the upper edge of the milling area and to one side of the target to enable drift correction over long acquisitions, and to allow a replacement fiducial to be milled if the fiducial score fell below 70%.

Serial milling was carried out with a 30 kV Ar⁺ beam at 60 pA. Block-face images were recorded in immersion mode with the through-lens detector (TLD) at 1.2 kV and 13 pA, with acquisition parameters optimized to minimize charging and rich cellular detail for each dataset. For *Anabaena inaequalis* (EMPIAR-14093) the TLD was operated in secondary electron mode (suction tube 70 V, mirror −15 V) with a 50 ns dwell time and 64-line integration, a slice thickness of 6 nm and a pixel size of 3 × 3 nm over a 10,500 × 6,300 nm field (3,500 × 2,100 pixels), giving a voxel size of 3 × 3 × 6 nm and an image time of 34.8 s per slice. For *Anabaena* sp. UTEX 2576 (EMPIAR-14092) the TLD was operated in down-hole visibility (DHV) mode (suction tube 244.7 V, mirror −15 V) with a 25 ns dwell time and 255-line integration, a slice thickness of 4 nm and a pixel size of 4 × 4 nm over a 5,248 × 5,248 nm field (1,312 × 1,312 pixels), giving an isotropic voxel size of 4 × 4 × 4 nm and an image time of 22.5 s per slice. Frame integration was 1 for both datasets. Complete acquisition parameters for both datasets are provided in supplementary **Table S1**.

### 2.5 Cryo-vEM segmentation

Annotation and segmentation were performed on a Hewlett Packard Z6 workstation operating on Windows 10 with 128 GB of RAM and a NVIDIA Quadro RTX 6000 with a 24GB GPU. TIF images were imported into Fiji (ImageJ, version 1.54t) and aligned using the built-in Scale-Invariant Feature Transform (SIFT) macro. The aligned images were imported into Dragonfly 3D World 2024.1 (Comet Technologies Canada Inc., Montreal, Canada) for segmentation. Next, the images were cropped to reduce stack size and exclude regions without sample, and a rolling convolution filter with a kernel size of 5 (**Figure S1**) (equivalent to the slab average feature) was applied using the Dragonfly 3D World image filtering function to reduce noise.

#### 2.5.1 Manual segmentation

The carboxysomes, cyanophycin granules, and cell walls were segmented manually in Dragonfly 3D World. A ROI was created for the cell wall, and the boundaries of the walls were segmented using the multi-slice pen. When the multi-slice pen is applied, segmentation on one image applies to the adjacent top and bottom 3 images for faster manual segmentation. Additional ROIs were created for the carboxysomes, cyanophycin granules and polyphosphate granules. Each was manually outlined and filled in using the “fill inner areas” function in the z-plane to enable volume measurements.

For 3D modeling (**Figure 1**, Image processing; **Figure 3E, F**) contour meshes for the cell wall and carboxysomes were created at 1:1:1 XYZ sampling. The opacity of the cell wall, cyanophycin granules, and carboxysome meshes were decreased for improved visualization of internal structures. The screenshot option was used in Dragonfly to export individual TIF images. Videos were created for each dataset using Dragonfly 3D World’s 2D animation and Movie Maker functions and exported as AVI files for *Anabaena inaequalis* (see **Video S1, S2**) and *Anabaena* sp. UTEX 2576 (see **Video S3, S4**). AVI files were converted to MP4 format with the H.264 video codec using FFmpeg (version 6.1.1; https://ffmpeg.org). Two-pass encoding was used to reach the target file size. The 2D videos were resized to 1280 × 818 pixels (**Video S2**) and 976 × 624 pixels (**Video S4**), with target bitrates of 1300 and 550 kbit/s, respectively. The 3D rendering videos were kept at their original resolution and compressed using target bitrates of 2400 kbit/s (**Video S1**) and 2350 kbit/s (**Video S3**). No frames were removed, and the image content was not otherwise altered. Video conversion was carried out with the assistance of Claude (Opus 5.5; Anthropic), and all output videos were checked by the authors.

**Figure 3.**
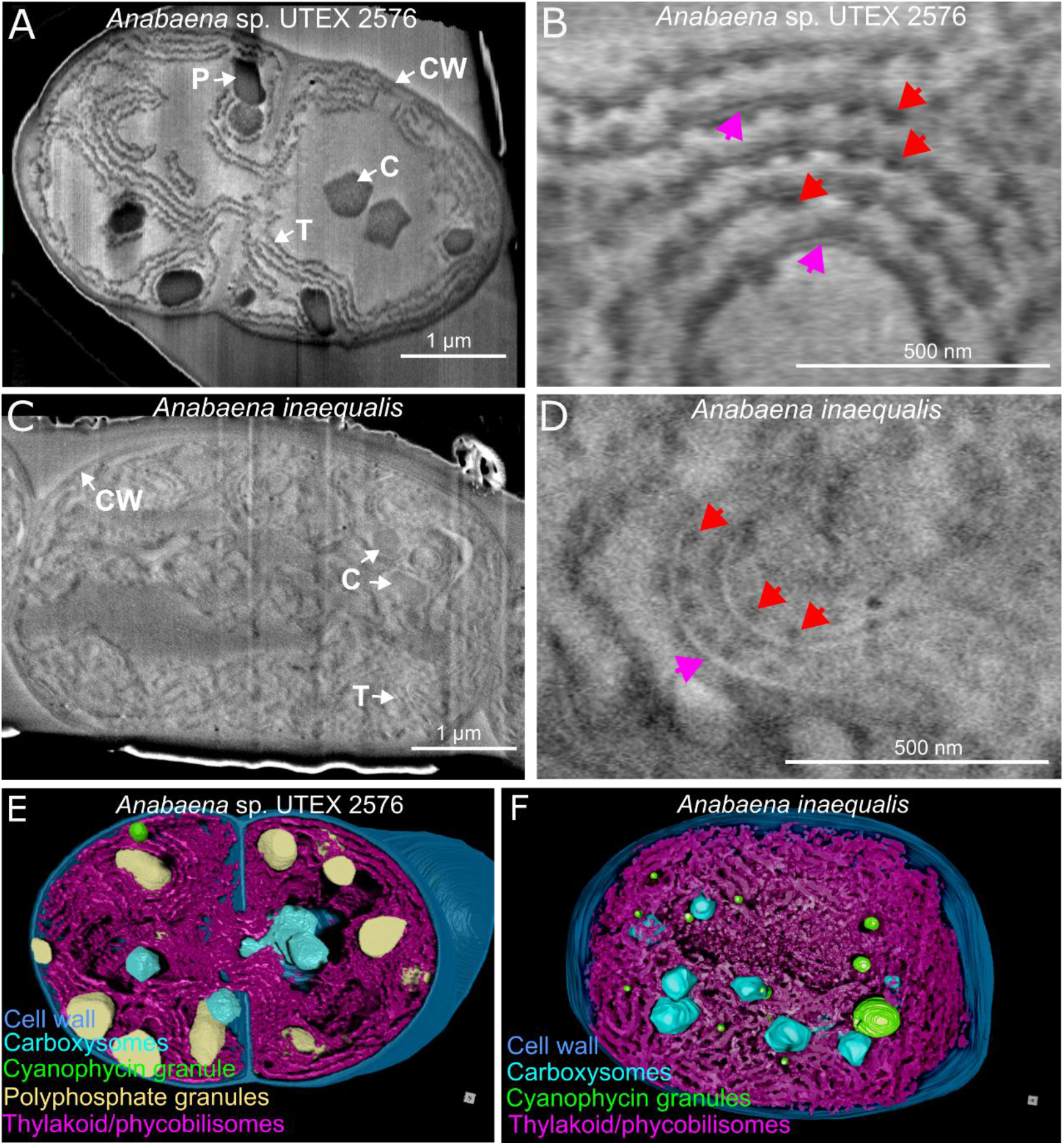
Cryo-vEM images and 3D modeling of two *Anabaena* species. (A) A single overview image from a 1300 slice cryo-pFIB-SEM z-stack of *Anabaena* sp. UTEX 2576 with contrasted cell wall (CW), thylakoid membranes with phycobilisomes (T), carboxysomes (C), and polyphosphate granules (P). (B) Higher magnification of the thylakoid double-membrane separated by lumen (magenta arrow) and decorated with amorphous phycobilisomes (red arrow) from *Anabaena* sp. UTEX 2576 (C) A single overview image from a 695 slice cryo-pFIB-SEM z-stack of *Anabaena inaequalis* showing the cell wall (CW), thylakoid membranes with phycobilisomes (T), and carboxysomes (C). (D) Higher magnification of the thylakoid double-membrane separated by lumen (magenta arrow) and decorated with amorphous phycobilisomes (red arrow) from *A. inaequalis* dataset. (E) 3D modeling of segmented carboxysomes, cyanophycin granules, polyphosphate granules, cell wall, and thylakoid membranes with phycobilisomes from *Anabaena* sp. UTEX 2576 done in Dragonfly 3D World. (F) 3D modeling of the carboxysomes, cyanophycin granules, cell wall, and thylakoid membranes with phycobilisomes from the *A. inaequalis* dataset.

#### 2.5.2 Deep learning segmentation of thylakoids/phycobilisomes and polyphosphate granules

Segmentation of the polyphosphate granules and thylakoids/phycobilisomes involved using Dragonfly 3D World’s (2024.1 and 2025.1) Pre-trained 2.5D (3 slices) U-Net Depth 5 model within the module Segmentation Wizard. Due to the challenge of reliably training the U-Net models to separate the thylakoids from the similarly contrasted and closely associated phycobilisomes, this option was not pursued. Each U-Net model shared the same parameters outlined in **Figure 4** (depth of 5, input slice count of 3, and patch size (pixels) of 64,64,1; See **Supplementary Thylakoids U-Net Model S1** and **S2** and **Supplementary Polyphosphate U-Net Model S3** on Zenodo at 10.5281/zenodo.23808628). In order to get rapid feedback, all models were initially trained by manually annotating 5 whole frames from each dataset with two classes: objects of interest and background. Once initial annotation of the training frames was complete, the model trained for a minimum of 10 epochs before it was tested by segmenting the objects of interest from the entire dataset and its success was evaluated by comparing segmentation output against the dataset. Areas where the model performed poorly in the segmentation were chosen as the focus for successive rounds of training to further improve output. Each additional round of training consisted of adding 5-8 new subframes of manual annotation in a new Segmentation Wizard session where the model was trained again for a minimum of 10 epochs. Following each additional round of training the model was again tested by segmenting the objects of interest from the entire dataset and results were evaluated. After the final round of training, the respective models were applied to the entire dataset and extraneous or missing pixels cleaned as needed from the segmentations.

**Figure 4.**
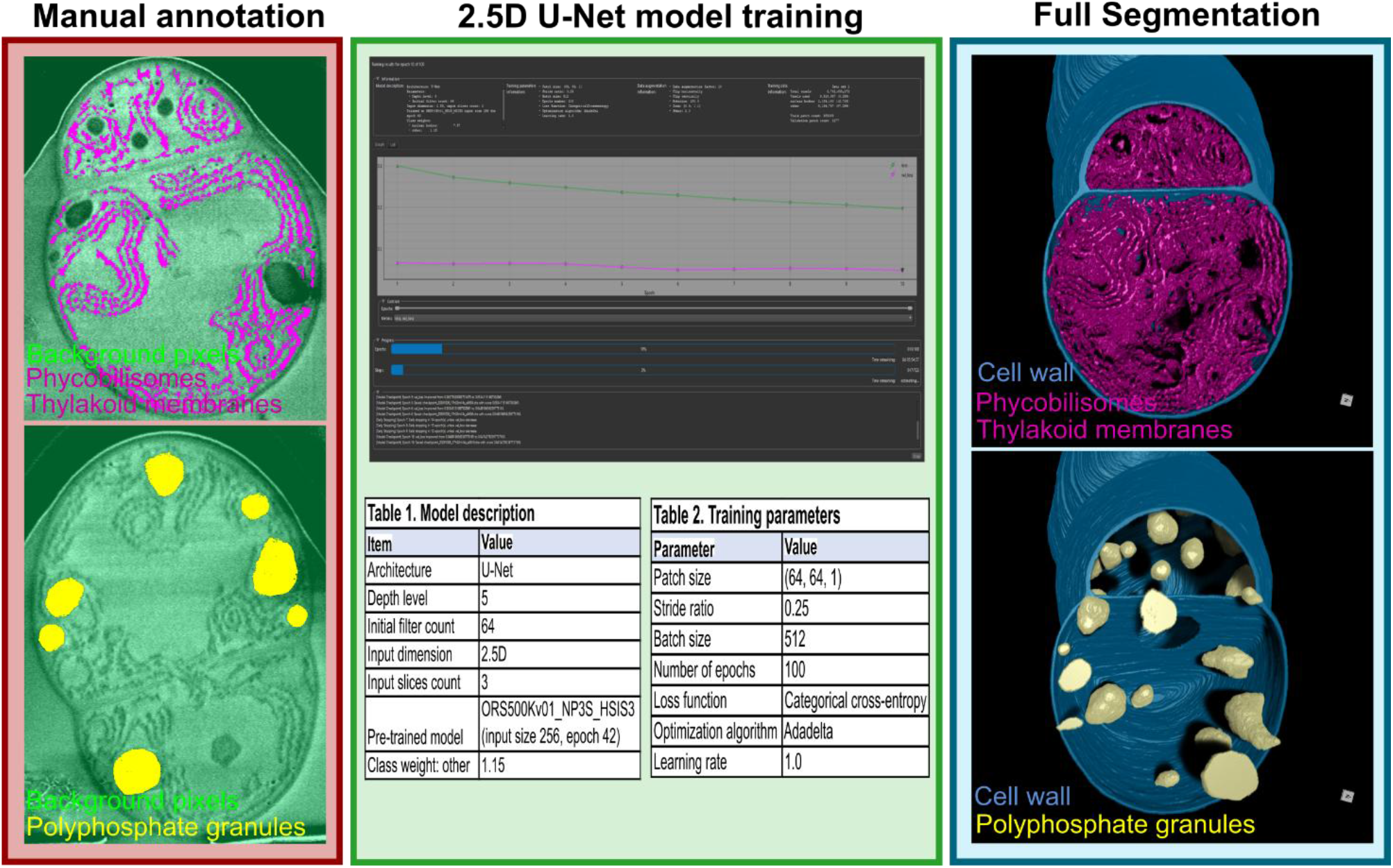
AI training workflow on *Anabaena* thylakoid membranes and polyphosphate granules using Dragonfly software. A training workflow for AI implementation: first, 4 – 5 whole image frames are manually annotated to train the U-Net AI. Then the model is trained for at least 10 epochs and up to 100 epochs as long as improvement in fit continues. Successive rounds of training can be applied to increase the specificity of the AI model. Once sufficiently trained, the AI model was applied to the entire dataset and cleaned as needed. 3D models displaying the thylakoids/phycobilisomes (magenta) and polyphosphate granules (yellow) after applying the trained AI model to the dataset along with the manually segmented cell wall (cyan) in *Anabaena* sp. UTEX 2576.

### 2.6 Quantification of carboxysomes and cyanophycin granules

After manual segmentation of carboxysomes and cyanophycin granules, they were converted to a “multi-ROI” in Dragonfly 3D World, separating each structure into an individual ROI for volume measurements (**Table S2**). Thylakoid double-membrane-plus-lumen thickness and the spacing between adjacent thylakoids were measured from individual block-face images using the line-measurement tool in Dragonfly 3D World (**Figure S2 and Table S3**). All measurements were exported as .csv files to continue working from Microsoft Excel. For paper submission, data tables were converted into Microsoft Word documents by Claude (Opus 5.5; Anthropic). The authors verified the data in the Microsoft Word document. The violin plots in **Figure 5** and histograms in **Supplementary Figure S3** were made using GraphPad Prism version 11.0.2.

**Figure 5.**
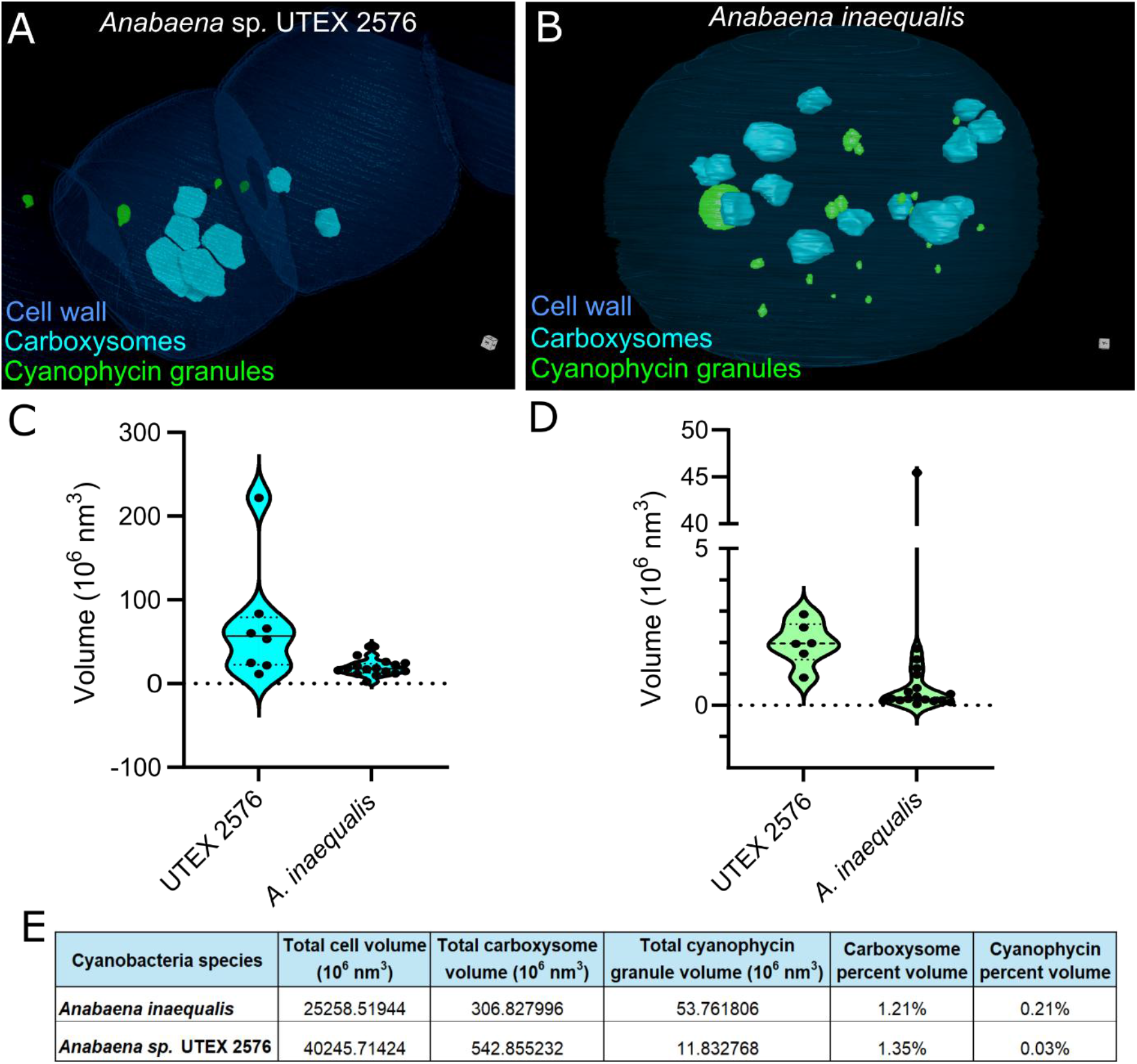
Quantification of carboxysomes and cyanophycin granules from cryo-vEM. (A) 3D modeling of the cell wall, carboxysomes and cyanophycin granules in *Anabaena* sp. UTEX 2576 showing their relative distributions within the cleaving cell. A small portion of the cleaving cell wall has not fully closed. (B) 3D modeling of an *Anabaena inaequalis* cell of the cell wall, carboxysomes, and cyanophycin granules. (C) Violin plot of comparative volume measurements of the carboxysomes (C), cyanophycin granules (D) between *Anabaena* sp. UTEX 2576 and *A. inaequalis*. (E) A table of total volume measurements and percent volumes of carboxysomes and cyanophycin granules. While individual carboxysome volume varied somewhat within the cells measured from these 2 species, carboxysome volume/cell volume was relatively constant while cyanophycin granules/cell volume showed a 7-fold difference.

## 3 Results

### 3.1 A cryo-vEM workflow for vitrified filamentous cyanobacteria

We established a generalized cryo-vEM workflow for *Anabaena* sp. that runs from grid preparation to quantified three-dimensional models (summarized in **Figure 1**) without chemical fixation, dehydration, heavy-metal staining or resin embedding at any step. Grids were rendered hydrophilic via glow discharge, treated with poly-L-lysine to improve filament attachment and loaded with culture in liquid media for 5 minutes, before vitrification via plunge-freezing into liquid ethane. Vitrified grids were clipped into AutoGrids and transferred to the cryo-pFIB-SEM, where they were mapped and individual filaments were located by a combination of secondary electron and/or integrated fluorescence light microscopy (iFLM) using the intrinsic autofluorescence of the pigments in photosynthetic cells. Next, the target frozen cells were protected with a platinum and xenon multilayer coating, trenched, and then imaged using automated argon milling and block-face detection using Auto Slice & View (AS&V). The resulting stacks were fine aligned, segmented, and quantified in Dragonfly 3D World. Two whole cell volumes were acquired, one per species: *Anabaena inaequalis* at 3 × 3 × 6 nm voxels and 695 slices (EMPIAR-14093) and *Anabaena* sp. UTEX 2576 at 4 × 4 × 4 nm isotropic voxels and 1312 slices (EMPIAR-14092), with the two filaments approached from different orientations — one from the longitudinally and one oblique cross-section to the filament axis — to test whether milling geometry constrained the result. Longitudinal sections of the filament in general require fewer slices, while oblique cross sections can run up a chain of cells and reduced image acquisition pixel dimensions for a faster per slice stack.

### 3.2 Poly-L-lysine retains filaments through blotting and vitrification

Reliably retaining filaments on the grid through blotting proved to be a somewhat limiting step in specimen preparation. Grids that were glow-discharged alone often carried only sparse, isolated filaments after blotting, with most grid squares empty (**Figure 2A**). Grids that were glow-discharged and then coated with poly-L-lysine retained a more dense, evenly distributed population of filaments spanning many contiguous grid squares (**Figure 2B**). The improvement persisted through vitrification: cryo-pFIB-SEM and iFLM maps of poly-L-lysine-treated grids showed filaments distributed across the grid within presumptive vitreous ice and suitable for targeting (**Figure 2C**). IFLM (**Figure 2D**) was very useful to efficiently confirm exact positioning and orientation of autofluorescent *Anabaena* cells beneath the thin ice surface, especially when they were clustered and/or did not protrude above the frozen film. **Figure 2E-G** documents key steps in the handling sequence, from incubation of the culture on the clamped grid, blotting against the filament applied face to storage of vitrified grids in the plunge-freezer cryo-chamber.

### 3.3 Native contrast enables cyanobacterial ultrastructure in both species

Single block-face images of vitrified filaments enabled visualization of the principal compartments of the cyanobacterial cell without any contrast agent. In *Anabaena* sp. UTEX 2576 the cell wall, concentric thylakoid membrane sheets, carboxysomes and electron-lucent storage inclusions were all directly visible in a single slice (**Figure 3A**), and the same inventory of features was contrasted in *Anabaena inaequalis* (**Figure 3C**). Carboxysomes appeared as sharply delimited polyhedral profiles, darker than the surrounding cytoplasm and readily differentiated from the thylakoid system. Thylakoid sheets typically ran as concentric layers following the contour of the cell and converging toward the poles and the septum. As expected for an unstained frozen-hydrated block face, contrast grey levels were inverted relative to heavy metal stained, resin-embedded FIB-SEM: the high water content cytoplasm appeared light and lipid-rich membranes dark, with contrast derived from the specimen’s composition. Closer inspection of *Anabaena* sp. UTEX 2576 (**Figure 3B**) and *Anabaena inaequalis* (**Figure 3D**) of thylakoid membranes cross-sections and phycobilosomes, revealed a thin double membrane and lumen as expected which were decorated by individual phycobilosome clusters. While some differences in relative contrast and image acquisition parameters were noted, overall image quality was comparable between the two datasets despite their different voxel sizes, beam/detector settings and milling orientations, species, and physiological states. This reassures us that the general conditions are somewhat versatile and not limited to one strain, approach geometry, and/or physiological state.

### 3.4 Segmentation yields complete three-dimensional models of single cells

Segmentation of the aligned stacks produced complete three-dimensional models of single vegetative cells in both species (**Figure 3E,F**). The cell wall, carboxysomes, cyanophycin granules were segmented manually, whereas the polyphosphate granules and thylakoid– phycobilisome compartment was segmented using a 2.5D U-Net trained on four to five manually annotated frames and then applied to the entire stack (**Figure 4**). Individual phycobilisomes were visible in the block-face images, but the model could not reliably assign them to a class separate from the membrane they decorate, so the two were segmented together as a single thylakoid–phycobilisome class. The reconstructions show this compartment as an interconnected system of sheets occupying most of the cell interior, with carboxysomes and cyanophycin granules asymmetrically distributed within and between the sheets rather than displacing them. Supplementary **Video S1** and **Video S2** show the segmented volumes and the underlying image stacks in z from *Anabaena inaequalis* and supplementary **Video S3** and **Video S4** from *Anabaena* sp. UTEX 2576, respectively.

### 3.5 Carboxysome and cyanophycin granule volumes differ between the two species

Because every inclusion was segmented in 3D, they could be counted and measured directly rather than inferred from single 2D sections. In the reconstructed *Anabaena* sp. UTEX 2576 cell, 8 carboxysomes were identified, with a mean volume of 67.9 ± 62.6 × 10^6^ nm^3^ (median 56.7, range 11.5–221.8 × 10^6^ nm^3^). The *Anabaena inaequalis* cell contained twice the number of carboxysomes, (16), but they were notably smaller and more uniform: mean volume 19.2 ± 9.6 × 10^6^ nm^3^ (median 17.7, range 2.3–44.1 × 10^6^ nm^3^). The distributions are shown as violin plots in **Figure 5C** and as histograms in **Supplementary Figure 3**. Despite this size and number difference, the carboxysome volume per cell was similar for these 2 species, at 542.8 × 10^6^ nm^3^ in UTEX 2576 and 306.8 × 10^6^ nm^3^ in *A. inaequalis*.

Cyanophycin granules showed a strikingly different pattern. The UTEX 2576 cell contained 6 granules of closely similar size, with a mean volume of 2.0 ± 0.6 × 10^6^ nm^3^ (range 0.88–2.9 × 10^6^ nm^3^), whereas the *A. inaequalis* cell contained 18 granules spanning three orders of magnitude, most of them very small (median 0.2 × 10^6^ nm^3^, range 0.035–45.4 × 10^6^ nm^3^) together with a single granule of 45.4 × 10^6^ nm^3^ (**Figure 5A, B, D**). Because the cell boundary itself was segmented, each inclusion class could be expressed as a fraction of the whole cell. The total cell volume was 25,259 × 10^6^ nm^3^ (25.3 µm^3^) for *A. inaequalis* and 40,246 × 10^6^ nm^3^ (40.2 µm^3^) for UTEX 2576. Carboxysomes had a similar proportion of each cell, 1.21 % in *A. inaequalis* and 1.35 % in UTEX 2576, despite the 2X difference in number and much different individual size. Cyanophycin, by contrast, differed more than 7X between the two cells, at 0.21 % and 0.03 % of cell volume, respectively (**Figure 5E**). Ultimately, the cryo-vEM approach proved to be highly appropriate to differentiate and quantify these important cyanobacteria sub-cellular features.

### 3.6 Direct measurement of thylakoid architecture in the vitrified state

Because the cells were imaged frozen-hydrated, without dehydration or resin infiltration, membrane spacings could be measured directly from the block-face images rather than inferred from otherwise chemically pre-treated and processed material. To evaluate these distances, we selected membranes oriented in cross-section (as opposed to oblique). In *Anabaena* sp. UTEX 2576 the thylakoid double membrane and lumen measured 14.5 ± 1.8 nm (mean ± SD, n = 5) and the spacing between adjacent thylakoid membrane pairs measured 76.2 ± 3.7 nm (n = 4) (**Supplementary Figure 2**). The paired membranes and the intervening lumen were directly resolved (**Figure 3B, D, magenta arrows**), and the stromal faces of the sheets were decorated with discrete phycobilisome densities (**Figure 3B, D, red arrows**), consistent with phycobilisomes occupying, and largely setting, the spacing between adjacent thylakoid sheets.

## 4 Discussion

Our initial challenge with cryo-vEM of *Anabaena* was not imaging itself, but retaining filaments on the grid in a usable density and distribution. With glow-discharged UltrAuFoil grids alone, few filaments were retained after blotting (**Figure 2A**), whereas grids additionally coated with poly-L-lysine retained a more even and dense population of filaments through blotting and vitrification (**Figure 2B**) that was still evident in the cryo-pFIB-SEM grid map (**Figure 2C**). Filamentous organisms are particularly prone here, because a filament spanning several grid squares presents far more surface for the blotting paper to directly contact than single cells do. The poly-L-lysine treatment was simple and included an overnight drying step to improve attachment properties, similar to coverslip functionalization used for light microscopy. Furthermore, we believe this straightforward approach be very useful for other cell systems when attempting cryo-vEM of filamentous, weakly/non adherent, or low concentration organisms.

A second important point is that *Anabaena* did not require high-pressure freezing. Vegetative cells while elongated in chains, as individuals are roughly 5 µm in diameter, well within plunge-freezing vitrification capabilities [5, 6], with the added benefit that no cryoprotectant or lift-out is needed. This is a meaningful simplification relative to high-pressure frozen (albeit, it does not preclude the use of HPF if desired), lift-out workflows that bulk tissue [38], and we believe cyanobacteria would be a highly accessible, and suitable entry point for laboratories beginning cryo-vEM of photosynthetic organisms.

Our quantification of (8 and 16) carboxysomes in a single vegetative cell are within the published range. For example, fluorescence green fluorescent protein labeled β-carboxysomes imaged by confocal microscopy in *Synechococcus elongatus* PCC 7942 give 1.8 ± 1.2 to 10.4 ± 3.8 per cell from low to high light [43], and whole-cell electron tomography of *Cyanothece* sp. ATCC 51142 resolved 21 in a single reconstructed cell [44]. Our mean diameter of 322 nm is larger than cryo-ET values for purified *Synechococcus* β-carboxysomes, which averaged around 193 nm over a 150–400 nm range [45] or 169.0 ± 11.8 nm [46], but our results are within the 250–600 nm range using electron tomography of *Cyanothece* [44] and larger than α-carboxysome values of approximately 125 nm [47, 48]. We are unaware of published β-carboxysome dimensions for a filamentous cyanobacterium, so we are unable to confirm if these differences reflect species, the intact versus purified state, developmental state, and/or growth conditions but are an exciting avenue of study where vEM resolution and quantification of other nanoscale cell features would be important.

We found no published estimate for carboxysome volume as a proportion of total cell volume in any cyanobacterium. Comparable whole-cell volume-fraction accounting has been reported in other single-celled organisms, for the organelles of a complete yeast cell by FIB-SEM [8]. The nearest comparator is thylakoid volume, measured at approximately 15 % of cell volume in *Cyanothece* by electron tomography of high-pressure frozen, freeze-substituted cells [44]. With 1.21 % of the cell being used for the carbon-fixing compartment is something that has interesting implications in studies of light intensity, heat treatments, CO₂ and nitrogen concentration, as well as other environmental conditions. The only prior whole-cell 3D study of a cyanobacterium, a computer-aided reconstruction of *Agmenellum quadruplicatum* from semi-thick serial sections of glutaraldehyde and osmium fixed, ethanol-dehydrated and resin-embedded cells, likewise reported counts with an average of six carboxysomes per cell (range 4–10) over more than 50 reconstructed cells, but no volumetric measurements [49]. This is presumably a consequence of its sectioning thickness: with serial sections of 0.25–1.0 µm, the slice interval approaches or exceeds the diameter of a carboxysome, so inclusions can be located and counted reliably but not measurable in3D. Near-isotropic voxels of a few nanometers via cryo-FIB-SEM used herein, removes that constraint.

In general, some cyanobacteria structures quantified here are prone to conventional chemical processing. Polyphosphate granules have been observed to poorly infiltrate and pop out of resin sections during sectioning, leaving voids [50], and the ultrastructure of cyanophycin granules is fixative-dependent[41, 51]. Interestingly, contrast and measurements of these structures readily could be made on our vitrified cells without any aldehyde, dehydration, resin or stain. The same rationale has recently applied on vitrified *Anabaena* sp. UTEX 2576 using synchrotron X-ray fluorescence of vitrified *Anabaena* sp. UTEX 2576 to minimize preparation artifacts [52], and offers a highly complementary elemental perspective of the same strain in our study.

A precedent exists for the benefits of cryogenic imaging at these scales with imaging whole frozen-hydrated *Synechococcus* cells. For example, Kaneko et al. measured a carboxysome shell of 5–6 nm versus the 3–4 nm reported from chemically fixed sections [53, 54]. While we were unable to reliable contrast the carboxysome shell, our thylakoid measurements of 14.5 ± 1.8 nm reported here is the full thickness of the thylakoid, both bounding membranes together with the lumen. Electron microscopy of high-pressure frozen freeze-substituted resin-embedded Synechocystis [55] reported the only cyanobacterial measurement we are aware of that includes the thylakoid membrane pair enclosing the lumen and measured at 15 nm, remarkably similar when vitrified and imaged here via vEM. For the spacing between adjacent thylakoids we measured 76.2 ± 3.7 nm in vitrified *Anabaena (UTEX)*, against 30–50 nm reported for freeze-substituted resin-embedded *Synechocystis* [56] and 50–60 nm for *Cyanothece* also prepared by high-pressure freezing, freeze-substitution and resin embedding [44]. The vitrified measurements for phycobilisome size from *Anabaena* sp. PCC 7120 are approximately 54 × 32 × 21 nm and thus projects ∼32 nm from the membrane surface [57]. Thus, two opposed antenna layers would account for at least 64 nm of that spacing, leaving little room to spare. Assuming no species or environmental factors influencing the spacing, these values may suggest that dehydration and/or resin infiltration and concomitant shrinkage impact the cytoplasmic spacing between thylakoid sheets (reducing) reflect chemical processing.

Our 2.5D U-Net readily separated cell wall, carboxysomes, and cyanophycin granules but could not be trained to separate thylakoid membrane from phycobilisome, and the two were therefore segmented as a single class (**Figure 4**). Notably, individual phycobilisomes were readily visible in the block-face images (**Figure 3B and 3D**). However, our model was unable to reliably assign a class separate from the membrane they decorate because the phycobilisome are of similar contrast and contiguous with the thylakoid surface [57]. Nevertheless, while not within the scope of the work here, we believe a method could be developed to quantify the number, position and density of the phycobilisomes within the entire 3D volume. Alternatively, cryo-ET of focused-ion-beam-milled *Synechocystis* does readily discriminate individual phycobilisomes and their stacked arrays within a lamella on the membrane [29], which is the rationale to leverage cryo-vEM for whole-cell intermediate resolution and, where desired, *in situ* cryo-ET connected workflows.

## 5 Conclusions

The primary strengths of the cryo-vEM approach are the speed from culture to image, the absence of any chemical pre-treatment, the capture of entire 3D nanoscale whole cells, and compatibility with correlative fluorescence and cryo-ET workflows. Photosynthetic organisms have a further advantage that is useful for this work, namely phycobilisome and chlorophyll autofluorescence allow cells to be easily located beneath the icy surface and mapped by cryogenic iFLM without labelling (**Figure 2D**), staining, or genetic modification that usually limits some correlative cryo workflows. When needed, the use of relevant vital exogenous stains or genetically encoded fusion proteins could allow even more precision for select targets. Despite these benefits, charging of unstained, poorly conducting frozen specimens remains a constraint on image quality [17], and the dose that vitrified cells can tolerate impacts the achievable signal-to-noise per slice. Due to the relatively small size of *Anabaena* the whole cell quantification reported here is relatively straightforward, and the carboxysome, cyanophycin granules and thylakoid nanoscale measurements reflect a demonstration of what is possible with our cryo-vEM workflow as opposed to population estimates using bulk measurements. The resolution of the Hydra microscope, voxel size employed (3-6 nm), and sample beam interactions limit what we are able to measure and interpret. Notably, particular features such as the carboxysome shell, photosystem structure, Rubisco packing and the septal junctions that connect cells within the filament were not distinct or better suited for cryo-ET of milled lamellae [30] and the combined thylakoid–phycobilisome merged U-Net models described above are another example of the limits we experienced.

Two directions follow from our method development work described here with our two test organisms. The first is biology. *Anabaena* differentiates into heterocysts under low nitrogen environments and moves metabolites between cells through septal junctions [39], and these processes are organized at the mesoscale that cryo-vEM is able to address. Imaging a vegetative cell–heterocyst boundary in the vitrified state, with carboxysome and cyanophycin distributions quantified on either side, and under varying nutrient or environmental conditions is an obvious experiment. The second is methodological. The workflow described here applies to the same instrument already used for whole-cell algal volumes [35], and the very same vitrified specimen can be carried forward into lamella preparation and cryo-ET, giving a multi-scale perspective of a single (or multiple) cell and is part of our ongoing work as we have applied to other systems [14].

## Acknowledgments

We acknowledge imaging support from the Advanced Bioimaging Laboratory (RRID:SCR_018951) and use of the Thermo Scientific Helios 5 Hydra CX DualBeam pFIB-SEM acquired through generous donor support to the Donald Danforth Plant Science Center.

## Research ethics

Not applicable.

## Informed consent

Not applicable.

## Author contributions

The authors have accepted responsibility for the entire content of this manuscript and approved its submission. Conceptualization: AA, KJC, TMBS, Specimen preparation: AA, KJC, LR, Cryo-vEM image acquisition: LR, Segmentation and quantification: AA, CL, Writing: AA, LR, KJC, TMBS, Supervision: KJC, TMBS, Funding acquisition: WC, KJC, TMBS

## Use of Large Language Models, AI and Machine Learning Tools

Deep learning segmentation was performed using the pre-trained 2.5D U-Net model within the Segmentation Wizard module of Dragonfly 3D World, as described in Section 2.5.2; model architecture, training parameters and training data are reported there and in Figure 4. Claude (Anthropic) was used during preparation of this manuscript for literature search and for draft language editing, and for checking internal consistency of figure citations, reference numbering and reported values. All AI-assisted text was reviewed and revised by the authors, who verified every cited reference against the primary source. The authors take full responsibility for the content of this manuscript.

## Research funding

This work was funded by the National Science Foundation PGR Award# 2433594 to T Burch-Smith, K Czymmek (Donald Danforth Plant Science Center) and W Chiu (Stanford).

## Data availability

The cryo-vEM volumes reported here have been deposited in EMPIAR under accession numbers EMPIAR-14093 (*Anabaena inaequalis*) and EMPIAR-14092 (*Anabaena* sp. UTEX 2576). The trained U-Net models have been deposited on Zenodo (10.5281/zenodo.23808628) and the measurement spreadsheets are provided as Supplementary Material (**Tables S2,S3**) [27].

## Supplementary Figures

**Supplementary Figure 1.**
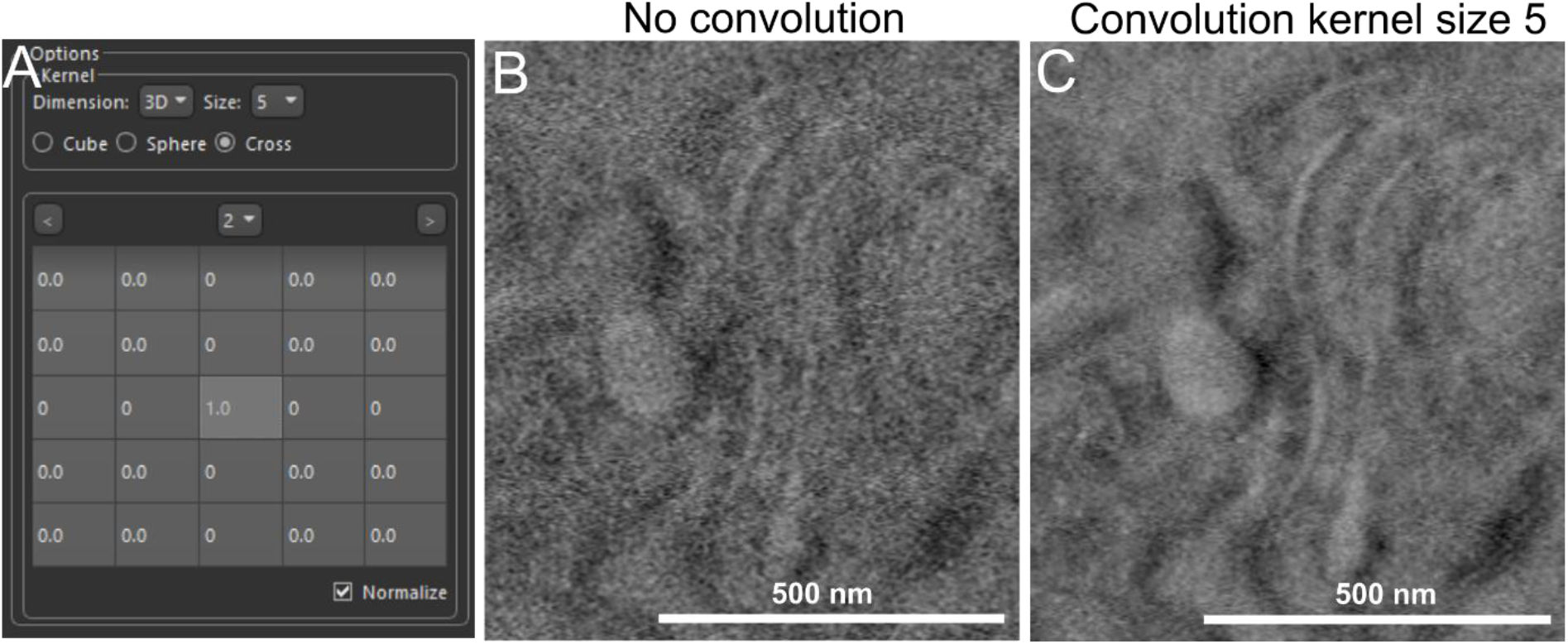
Method of convolution applied to *Anabaena inaequalis* images. (A) In Dragonfly 3D World, a convolution image filter was applied to the *A. inaequalis* images. The convolution filter was originally in a cross pattern, but the arms of the cross pattern were modified to “0” rather than “1”. The convolution was applied with a kernel size of 5 in the 3D dimension, this is equivalent to a 5-image slab average. (B) A zoomed-in image from *A. inaequalis* prior to applying the convolution filter. (C) An image from the same location as (B) after the convolution filter was applied.

**Supplementary Figure 2.**
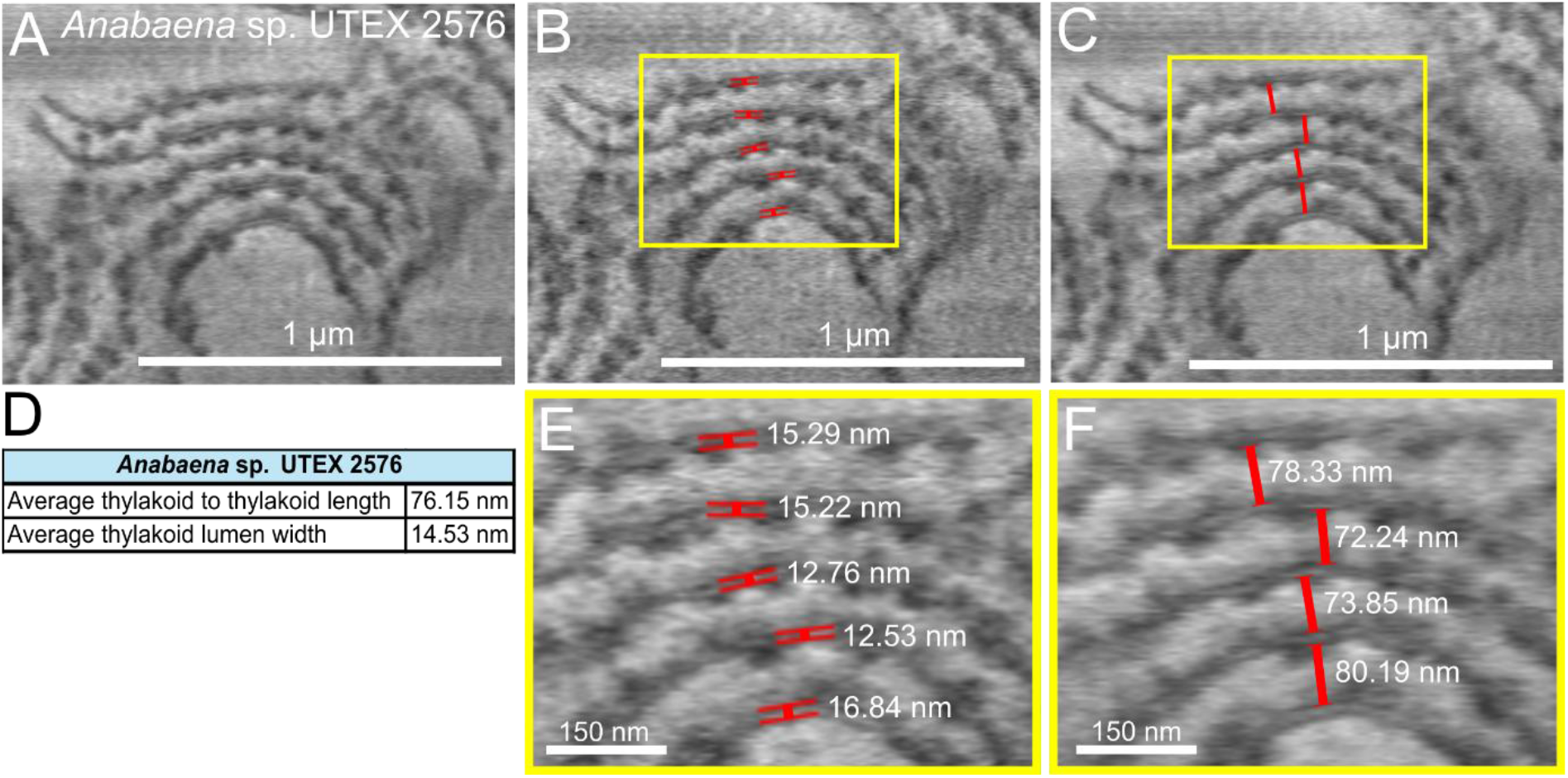
Quantitative analysis of thylakoid membranes in *Anabaena* sp. UTEX 2576. (A) A zoomed-in image of several thylakoid membranes. (B) A representative image showing the region of thylakoid membranes measured for double membrane and lumen width. (C) An image showing the region of measuring the distance between thylakoid membranes. (D) A table displaying the average measurement of thylakoid-to-thylakoid distance and the average thylakoid lumen width. (E) A zoomed-in image showing the thylakoid lumen measurements. (F) A zoomed-in image showing the measurements of the distance between the thylakoid membranes.

**Supplementary Figure 3.**
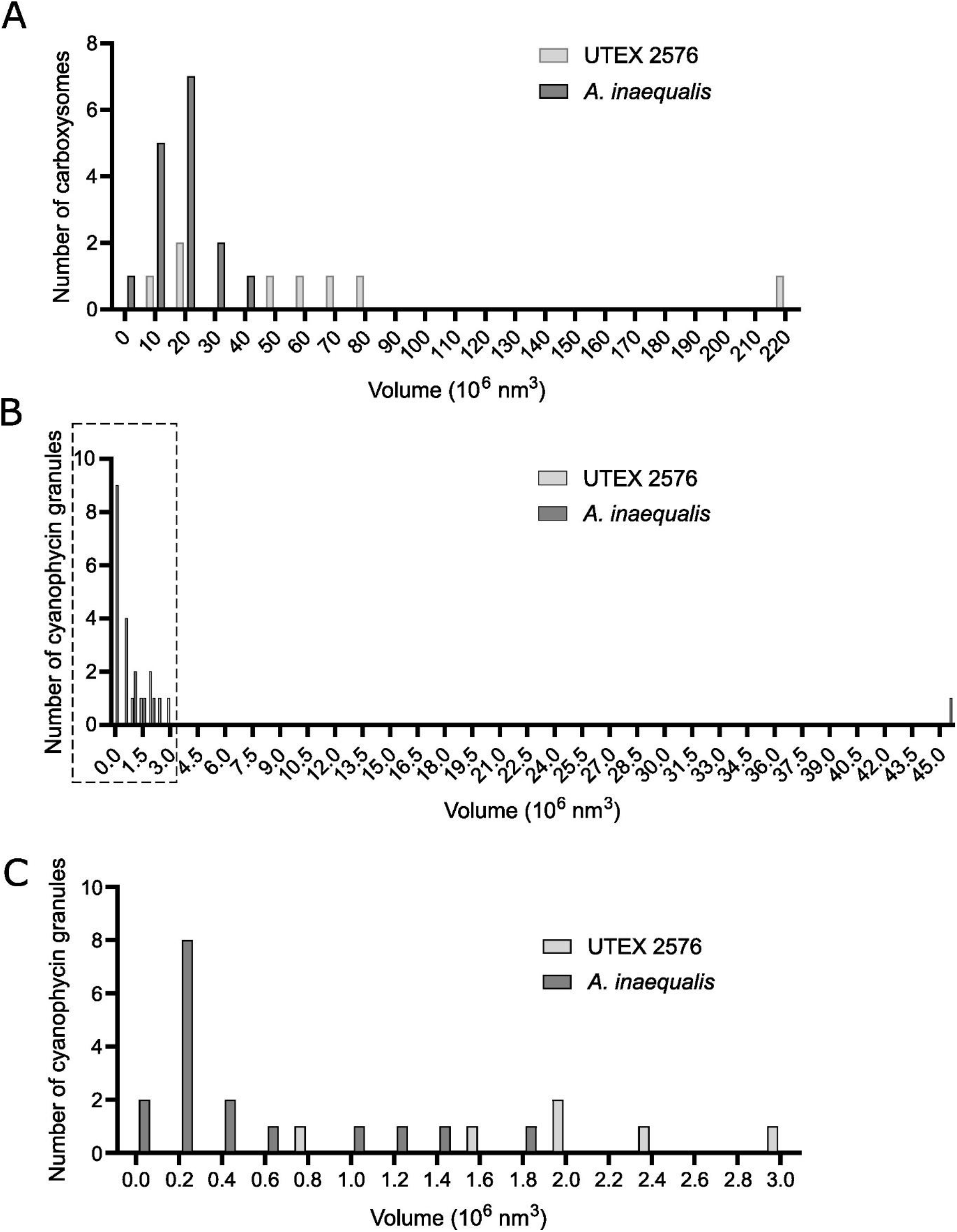
Histograms of carboxysome and cyanophycin granule volume measurements from cryo-vEM. (A) Histogram showing the distribution of carboxysomes volumes in *A. inaequalis* and *Anabaena* sp. UTEX 2576. (B) A histogram showing the volume distribution of the cyanophycin granules in both *Anabaena* species. Gray dotted box indicates the histogram in (C). (C) A zoomed-in histogram from (B) to display the majority of cyanophycin granule volumes.

**Table S1.**
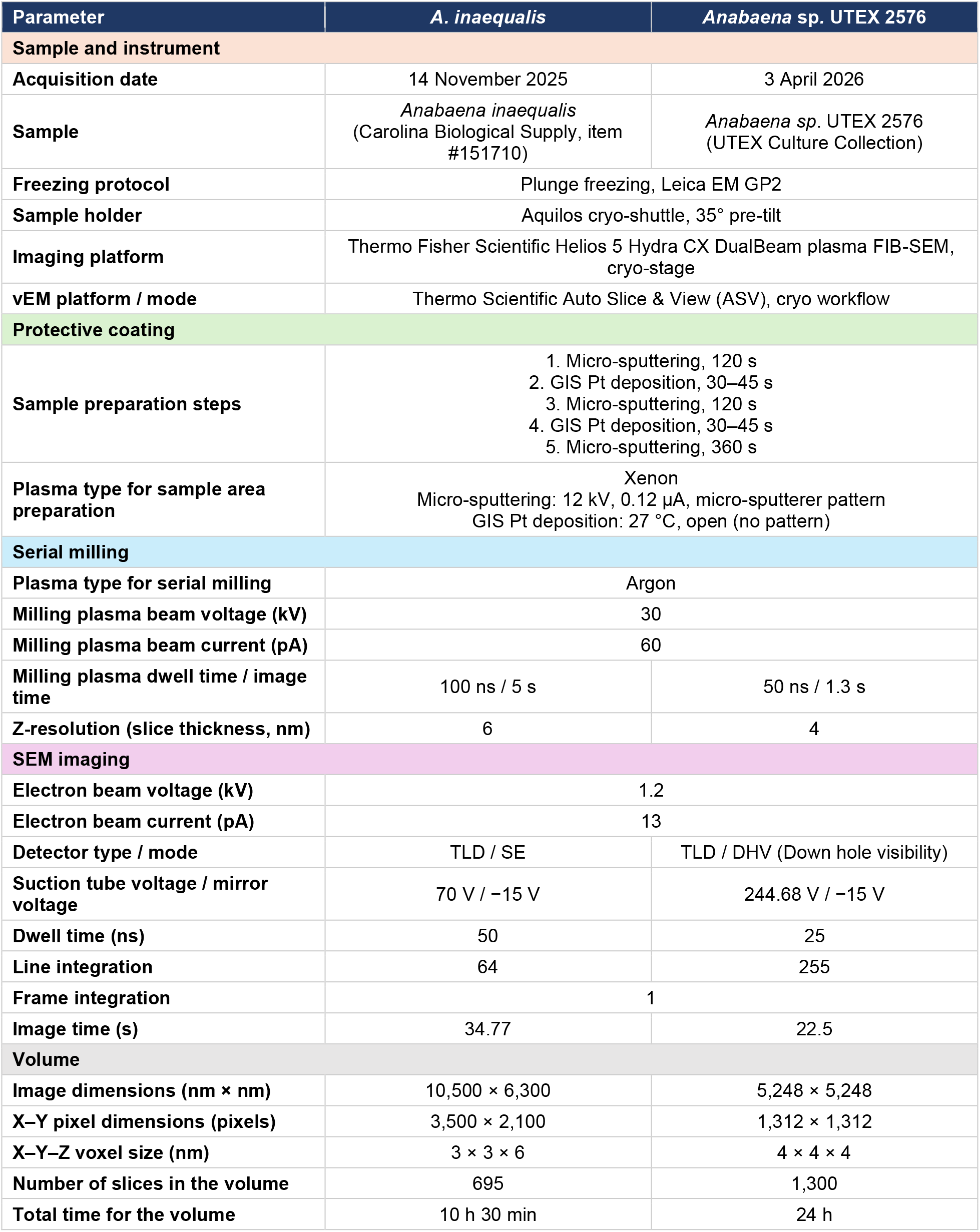
Cryo-pFIB-SEM acquisition parameters for whole cell volumes in *A. inaequalis* and *Anabaena sp.* UTEX 2576.

**Table S2.** Carboxysome, cyanophycin granule, and whole cell volumes in *A. inaequalis* and *Anabaena* sp. UTEX 2576.

| Carboxysome volume (10<sup>6</sup> nm<sup>3</sup>) |  |  |
|---|---|---|
| Carboxysome | ***A. inaequalis*** | ***Anabaena*** **sp. UTEX 2576** |
| 1 | 9.70 | 11.54 |
| 2 | 11.35 | 21.87 |
| 3 | 12.07 | 24.81 |
| 4 | 14.52 | 53.09 |
| 5 | 15.97 | 83.53 |
| 6 | 14.92 | 221.80 |
| 7 | 17.29 | 65.91 |
| 8 | 18.16 | 60.30 |
| 9 | 18.56 | – |
| 10 | 20.75 | – |
| 11 | 22.61 | – |
| 12 | 26.07 | – |
| 13 | 33.96 | – |
| 14 | 44.12 | – |
| 15 | 24.43 | – |
| 16 | 2.32 | – |
| **Average** | **19.18** | **67.86** |
| **Median** | **17.72** | **56.69** |
| **Cyanophycin granule volume (10<sup>6</sup> nm<sup>3</sup>)** |  |  |
| **Granule** | ***A. inaequalis*** | ***Anabaena*** **sp. UTEX 2576** |
| 1 | 0.549 | 1.956 |
| 2 | 0.035 | 2.477 |
| 3 | 0.096 | 1.976 |
| 4 | 0.131 | 0.882 |
| 5 | 0.158 | 2.901 |
| 6 | 0.198 | 1.642 |
| 7 | 0.974 | – |
| 8 | 0.179 | – |
| 9 | 45.438 | – |
| 10 | 0.128 | – |
| 11 | 1.183 | – |
| 12 | 0.358 | – |
| 13 | 1.464 | – |
| 14 | 0.211 | – |
| 15 | 0.277 | – |
| 16 | 1.801 | – |
| 17 | 0.429 | – |
| 18 | 0.153 | – |
| **Average** | **2.987** | **1.972** |
| **Median** | **0.244** | **1.966** |
| **Whole cell volume (10<sup>6</sup> nm<sup>3</sup>)** |  |  |
|  | ***A. inaequalis*** | ***Anabaena*** **sp. UTEX 2576** |
| **Whole cell** | 25,258.52 | 40,245.71 |

**Table S3.** Thylakoid measurements in *Anabaena* sp. UTEX 2576.

| **Thylakoid measurements (*Anabaena* sp. UTEX 2576)** |  |  |
|---|---|---|
| **Measurement** | **Thylakoid-to-thylakoid spacing (nm)** | **Thylakoid lumen width (nm)** |
| 1 | 78.33 | 15.29 |
| 2 | 72.24 | 15.22 |
| 3 | 73.85 | 12.76 |
| 4 | 80.19 | 12.53 |
| 5 | – | 16.84 |
| **Average** | **76.15** | **14.53** |
| **Median** | **76.09** | **15.22** |

